# Bootstrapping promotes the RSFC-behavior associations: an application of individual cognitive traits prediction

**DOI:** 10.1101/800243

**Authors:** Lijiang Wei, Bin Jing, Haiyun Li

## Abstract

Resting state functional connectivity records enormous functional interaction information between any pair of brain nodes, which enriches the prediction of individual phenotypes. To reduce the high dimensional features in prediction, correlation analysis is a common way for feature selection. However, rs-fMRI signal exhibits typically low signal-to-noise ratio and correlation analysis is sensitive to outliers and data distribution, which may bring unstable and uninformative features to subsequent prediction. To alleviate this problem, a bootstrapping-based feature selection framework was proposed and applied on three widely used regression models: connectome-based predictive model (CPM), support vector regression (SVR) and least absolute shrinkage and selection operator (LASSO). A large open-source dataset from Human Connectome Project (HCP) was adopted in the study and a series of cognitive traits were acted as the prediction targets. To systematically investigate the influences of different parameter settings on the bootstrapping-based framework, a total of 216 parameter combinations were evaluated through the R value between the predicted and real cognitive traits, and the best identified performance among them was chosen out as the final prediction accuracy for each cognitive trait. By using bootstrapping without replacement, the best performances of CPM with positive and negative feature sets, SVR and LASSO averagely increased by 28.0%, 33.2%, 11.6% and 24.3% in R values in contrast to the baseline method without bootstrapping. By using bootstrapping with replacement, these best performances increased by 22.1%, 22.9%, 9.4% and 19.6%. Furthermore, the bootstrapping-based feature selection methods could effectively refine the original feature sets obtained from correlation analysis, which thus retained the more stable and informative feature sets. The results demonstrate that bootstrapping-based feature selection is an easy-to-use and effective method to improve RSFC prediction of cognitive traits and is highly recommended in future RSFC prediction studies.

## 1 INTRODUCTION

Resting state functional connectivity (RSFC) has been found to be associated with pathological conditions and physiological status of human brain (Cribben et al., 2012; Elanbari et al., 2014; Finn et al., 2015; Meskaldji et al., 2016). Decoding RSFC provides an insight into understanding the neural mechanism underlying intrinsic brain activities (R. T. Jiang et al., 2018). Previous studies have reported that FC patterns could be applied to predict individual behavioral and cognitive phenotypes, such as attention ability (Yoo et al., 2018), cognition (Cui et al., 2018; Kong et al., 2019; Sun et al., 2019), intelligence ability (Finn et al., 2015; R. Jiang et al., 2019; Santarnecchi et al., 2014) and chronological age (Dosenbach et al., 2010; Liem et al., 2017). To model the RSFC-phenotype association, three prediction models including connectome-based predictive modeling (CPM), support vector regression (SVR) and least absolute shrinkage and selection operator (LASSO) have been frequently adopted for their good performances and interpretabilities (Coloigner et al., 2016; de Vos et al., 2018; R. Jiang et al., 2019; Meng et al., 2017; Ryali et al., 2012; Shen et al., 2017). CPM is in fact a linear regression model and has been successfully applied in predicting individual cognition (Finn et al., 2015; R. Jiang et al., 2019; Sripada et al., 2019), attention (Rosenberg et al., 2017), cocaine abstinence (Yip et al., 2019) and reading comprehension (Jangraw et al., 2018). SVR implements space conversion by some kernel function in order to achieve better prediction in new data space than in original data space (Basak et al., 2007), and has been utilized in prediction of mental disease (Rizk-Jackson et al., 2011), brain maturity (Dosenbach et al., 2010; Nielsen et al., 2019) and painful sensation (Tu et al., 2015). LASSO provides sparse representation by driving redundant features to zero-valued weights, and performs well in investigation of reward-related behavior (Ferenczi et al., 2016), mental state (Haufe et al., 2014) and various aspects of cognition (R. Jiang et al., 2019; R. T. Jiang et al., 2018; Rashid et al., 2014; Toiviainen et al., 2014).

For the high-dimensional RSFC, feature reduction is indispensable since redundant or irrelevant information may confound the statistical testing significance (Bunea et al., 2011), worsen the machine learning model performance (Arbabshirani et al., 2017; Duangsoithong et al., 2010) and increase computational complexity. For the regression tasks, correlation analysis between RSFC and the target phonotypic measure (e.g. cognitive trait) is the most common approach to select features (Dadi et al., 2019; Gabrieli et al., 2015). However, correlation analysis may not be able to catch the reliable correlational relationship since rs-fMRI data exhibits low signal-to-noise ratio (SNR) and correlation analysis is sensitive to outliers (R. Wilcox, 2004; R. R. Wilcox, 2005). Small perturbation in the data may lead to inclusion of unrelated features or exclusion of useful features, thus the weak robustness of correlation can introduce false correlations including type I error and power problem (Rousselet et al., 2012). In addition, data distribution may be another easily ignored factor in the correlation analysis: it is also susceptible to the clustered points, curvature, heteroscedasticity and range (Rousselet et al., 2012). All these conditions may cause that the features detected from the whole dataset did not significantly correlate with the target phonotypic measure in a subset, resulting in unstable features, which may in fact contribute little to the target phonotype prediction. Thus, an adequate feature selection method that could detect intrinsic stable RSFC features is of great importance to the prediction task.

Bootstrapping is a statistical method that relies on the random sampling (Efron et al., 1997; Hall et al., 2009), and any focused statistical test can be examined in newly resampled datasets. After repeating the resampling process several times, the stability of the statistical significance in each resampled dataset could be accessed, which could be finally used to ascertain the statistical test by setting specific threshold. The main advantage of bootstrapping is that it does not rely on any assumption on the data, so it can ascertain the actual and stable between-group statistical significance. A few literatures have adopted bootstrapping to investigate the stability of selected features. In Alonso-Atienza et al. (2012), bootstrapping was used as a backward feature selection method for cardiac ventricular fibrillation discrimination. Ditzler reported that bootstrapping combined with Neyman-Pearson hypothesis test successfully detected the statistically relevant features on both synthetic and real data (Ditzler et al., 2015). In Bunea et al. (2011), features which were selected most frequently by penalized least squares regression methods in bootstrapping resamples were identified as useful predictors in neuroimaging. However, few studies tried to explore the influence of different parameter settings on the bootstrapping. For example, Bunea et al. (2011) used bootstrapping with only one cutoff threshold (50% inclusion frequency) to derive the features. Abram et al. (2016) proposed a quantile threshold bootstrapping method for feature selection in penalized regression models. Hence, studies were highly needed to make known how different parameter settings in bootstrapping influenced the final prediction performance.

This study explored a bootstrapping-based feature selection framework to enhance the RSFC prediction performance in CPM, SVR and LASSO models, and different parameter settings in bootstrapping were specially compared in order to ascertain the optimal parameter setting, which mainly included two categories: bootstrapping without replacement (i.e., subsample without replacement) and bootstrapping with replacement. To verify these bootstrapping-based feature selection methods, a large public dataset from Human Connectome Project was used in this study, and 13 cognitive traits were selected as the prediction targets. We speculated that bootstrapping could improve the RSFC prediction performances of CPM, SVR and LASSO models.

## 2 METHODS AND MATERIALS

### 2.1 Resting-state fMRI data

The data came from publicly WU-Human Connectome Project (HCP) 1200 release. All imaging data were collected on a 3T customized Siemens scanner using a standard 32-channel head coil with a multiband pulse sequence. Two scanning sessions (REST1 and REST2) of high spatial-temporal resolution resting state fMRI (rs-fMRI) data were acquired during two consecutive days. Both sessions were scanned at right-to-left and left-to-right phase encoding direction, and the functional images comprised 1200 volumes in one session. For more details of inclusion and exclusion criteria on the dataset, please see Van Essen et al. (2013).

### 2.2 Preprocessing

The extensively preprocessed PTN (Parcellation+ Timeseries+ Netmats) data were utilized in our study. Details of preprocessing steps could be found in Smith et al. (2013). In brief, each run of rs-fMRI data underwent a minimal preprocessed pipeline and ICA+FIX to remove the potential artifacts. All 4 runs were concatenated to form 4800 volumes for each individual. After region parcellation by group-ICA, the subject-specific FC matrices, or connectomes, were calculated using Pearson’s correlation and then Gaussianised into Z-stats. Here, we adopted 300 ICA components as network nodes, resulting in a connectivity matrix with size 300×300 for each participant. Notably, there were two types of fMRI data reconstruction methods available in the release, and it was not clear the potential influences induced by different reconstruction algorithms. To avoid the heterogeneity in data, the current study limited the analyses to 812 healthy subjects (ages 22-37) whose rs-fMRI data were reconstructed using a later improved version of construction method, and discarded data with original reconstruction version.

### 2.3 Cognitive traits

Given that the cognitive activities were functionally modulated by the brain, prediction of cognitive traits by RSFC was a feasible task (Stevens, 2009). Previous studies had shown that RSFC patterns were closely associated with cognitive traits (Beaty et al., 2018; Sun et al., 2019). In this study, 13 measures of cognitional tests were chosen as the prediction targets that provided by HCP (Li et al., 2019). These tests included Picture Sequence Memory, Dimensional Change Card Sort, Flanker Task, Oral Reading Recognition, Picture Vocabulary, Pattern Completion Processing Speed, List Sorting, Penn Progressive Matrices, Delay Discounting, Variable Short Penn Line Orientation, Short Penn Continuous Performance and Penn Word Memory. Seven of them measured by NIH (National Institutes of Health) Toolbox were normalized to age-adjusted scores with mean of 100 and standard deviation of 15. Furthermore, not all subjects had the whole 13 cognitive scores and subjects with missing cognitive data were excluded in subsequent analysis.

### 2.4 Bootstrapping-based feature selection methods

The bootstrapping-based feature selection methods sought to find stable features that were consistently identified in the resampling subsets. Conventional bootstrapping method mainly focused on resample with replacement, but it usually required a high amount of resample times, which seemed not efficient enough for feature selection. Moreover, medical imaging dataset was usually not large enough, in this condition, bootstrapping without replacement might suit the finite population. In this study, a wide range of parameter settings on bootstrapping with and without replacement were tested and evaluated to make sure the optimal parameter setting of bootstrapping.

#### 2.4.1 Bootstrapping without replacement

Bootstrapping without replacement took out samples without replacement from the original dataset, so each subject in the resampling dataset was unique without duplication. For each cognitive measure, the bootstrapping without replacement was applied on training set to extract the feature vector. Let *NB* (number of bootstrapping) denote the sampling times. Within each resampling iteration, a portion of subjects were randomly selected without repetition. The proportion of sampling was defined as bootstrap percentage (*BP*), e.g., *BP* = 50% indicated half of all subjects were chosen from original dataset to form a resample subset. Spearman correlation analysis was then conducted in each resampling set between RSFC and cognitive measure, and significant features were selected as predictors. After all bootstrapping iterations finished, a stability threshold for frequency percentage (*FP*) of feature was used to determine final feature sets in *NB* bootstrapping datasets. For example, if the stability threshold was set to 50%, then feature whose *FP* more than 50% would be selected in the final feature set, otherwise it would be filtered as a weak predictor.

#### 2.4.2 Bootstrapping with replacement

In bootstrapping with replacement method, resamples were obtained with replacement from original samples, so the sample size of resampling subset was same as original dataset size. There were only two parameters in this method, i.e. *NB* and *FP*. After performing Spearman correlation analysis in *NB* bootstrapping subsets, features with *FP* larger than stability threshold were chosen as predictors.

### 2.5 Regression models

Three widely used regression models were employed in the study to predict the cognitive traits with RSFC, including CPM, SVR and LASSO.

#### 2.5.1 CPM

Briefly, CPM summarized the relevant RSFCs for each individual and fitted the summary values with cognitive measures in a linear regression model. The detail steps of CPM were as follows:

1. Find features in RSFC matrices that correlated with the cognitive measure, and significantly positive or negative features were respectively selected.
2. Summarize selected feature values of positive feature set and negative feature set separately for each individual.
3. Perform univariate linear regression between the summary value and the cognitive measure for positive and negative feature sets separately.
4. Test the model performance with the unseen testing data.

Here, label the CPM model using positive or negative feature sets respectively as CPM-P or CPM-N.

#### 2.5.2 SVR

SVR was based on statistical learning theory and fitted linear model with Vapink’s ε-sensitive loss function (Smola et al., 2004). It tried to solve an objective function f(x) whose predicted values of training data deviated by no more than ε from actual values and flatness of regression line was maximized. Samples that deviated by more than ε from their actual values were called support vectors. Given N training samples ((x_l_,y_l_),…,(x_N_,y_N_)), the objective function took the following form:

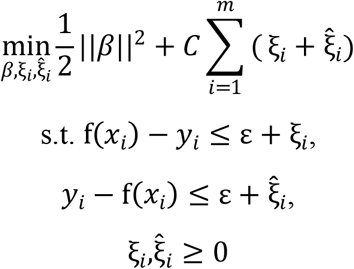

where *β* was the regression coefficient vector for features, *m* was the number of support vectors, ξ_i_ and 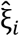 were slack variables and *C* was the penalty parameter that controlled the trade-off of penalty between variance and bias. The objective function was solved to obtain a weight for each support vector. In this study, the linear kernel was selected for SVR.

#### 2.5.3 Lasso

Lasso was a penalized regression method by adding L1-norm regularization to ordinary least squares (Tibshirani, 1996). The method could attenuate regression coefficients of correlated features except one feature among them to zeros (Zou et al., 2005), thus generated robust coefficients as well as parsimonious models. This algorithm minimized ordinary least squares plus the sum of absolute values of regression coefficients *β* to obtain the estimated coefficients. The objective function was formed as below:

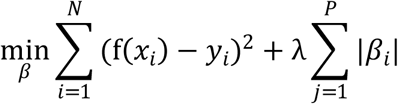

where *λ* was the tuning parameter which controlled balance between accuracy and sparsity.

At last, CPM, SVR and LASSO accompanied with correlation analysis on the whole dataset were marked as the baseline method, and the corresponding models combined with the bootstrapping-based feature selection were regarded as the proposed method.

### 2.6 Prediction Framework

For each cognitive score, a 10-fold cross-validation (CV) framework was utilized in the study. All individuals were sorted according to values of cognitive measure and then divided into 10 folds. To be specific, the (1^st^, 11^st^, 21^st^ …) subjects were assigned to 1^st^ fold, the (2^ed^, 12^ed^, 22^ed^ …) subjects were assigned to 2^ed^ fold and so on. This partition procedure ensured the same distribution of 10 folds and avoided random bias and expensive computation due to random splitting (Cui et al., 2018). Within the prediction framework, each fold was iteratively used for testing while the remaining 9 folds were used for training. The bootstrapping feature selection methods were performed on training set to select features for CPM, SVR and LASSO models (Figure 1). Significantly positive-, negative- and combined-correlated FCs were separated into positive, negative and combined feature sets (significant threshold p=0.05), which were respectively used to build CPM-P, CPM-N, SVR and LASSO models. In both bootstrapping method with and without replacement, a broad set of parameters including *NB*, *BP* and *FP* were tested, where *NB* was chosen from [10, 20, 50, 100, 500, 1000] and *FP* from [50%, 60%, 70%, 80%, 90%, 100%]. Besides, in bootstrapping without replacement, *BP* was chosen from [25%, 50%, 60%, 70%, 80%]. All parameter settings resulted in 180 combinations in bootstrapping without replacement and 36 combinations in bootstrapping with replacement. For SVR, an inner 5 fold-CV was employed to determine optimal parameters *C*, choosing from [10^−10^, 10^−9^… 10^10^]. For LASSO, an inner 10 fold-CV was employed to determine the optimal *λ* from range l0^−4^ to 1 with step size l0^−2^. The model performance was evaluated by the *R* value between predicted and actual cognitive values. In the study, the reported *R* values were averaged over 10 testing folds. All procedures were carried out with a multi-core CPU (Intel(R) Xeon(R) E5-2630 v4 @2.20GHz, 10 cores, 20 threads) with 128 GB memory.

**Figure 1.**
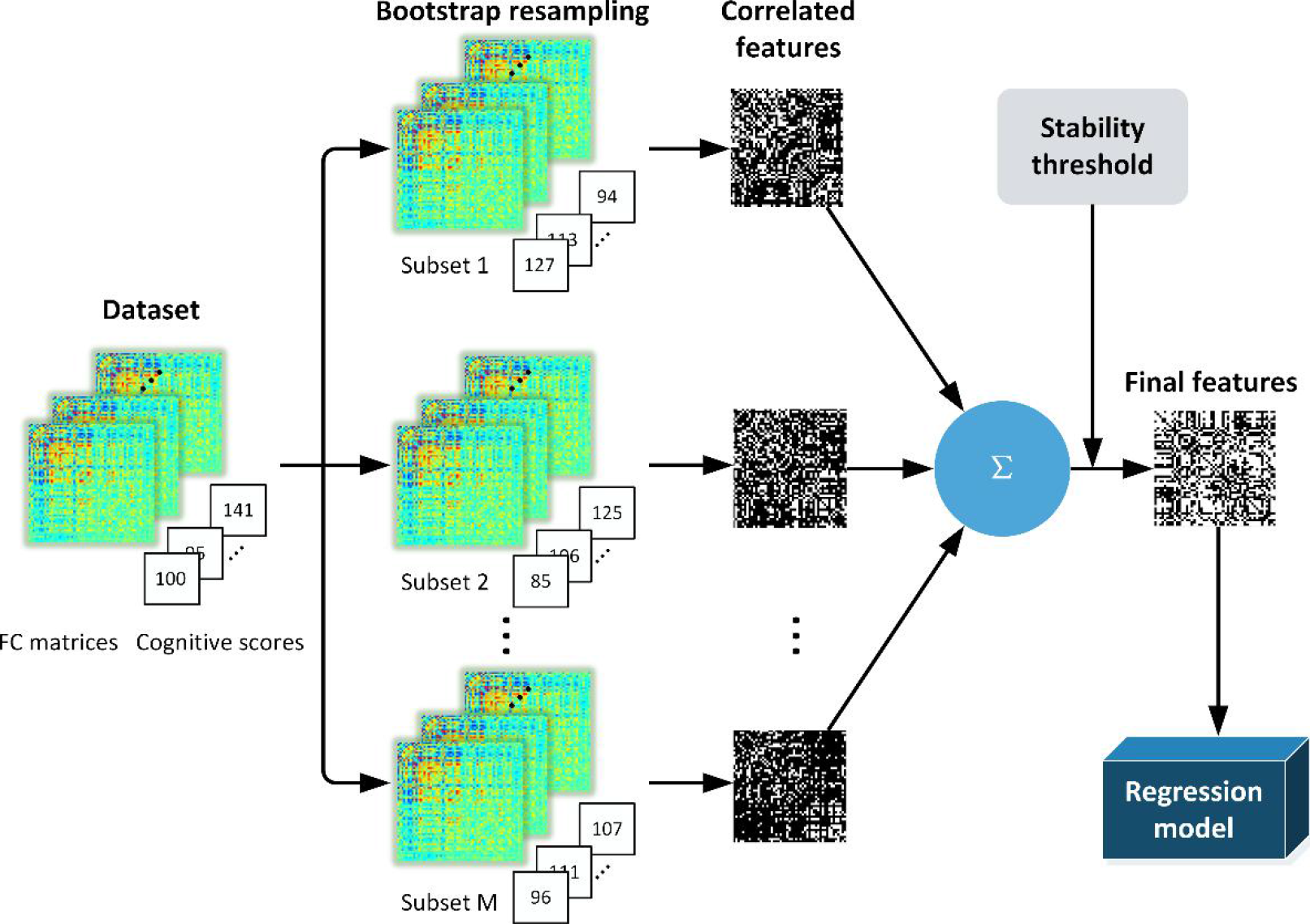
The pipeline of bootstrapping framework.

## 3 RESULTS

### 3.1 Bootstrapping improved prediction of cognitive traits

Figure 2 summarized the best prediction accuracies of 13 cognitive traits prediction achieved by bootstrapping methods and baseline method. All models using the refined feature sets derived from bootstrapping method outperformed the baseline method. Compared with baseline method (average r=0.15, 0.15, 0.24 and 0.18 for CPM-P, CPM-N, SVR and LASSO separately), the bootstrapping without replacement method increased the predictive correlation values by 28.0%, 33.2%, 11.6% and 24.3%. Among four models, SVR performed best and reached an average r=0.27, LASSO achieved suboptimal performance at average r=0.23 and CPM-P and CPM-N achieved similar performances of average r= 0.19 and r=0.20. In bootstrapping with replacement, the accuracy increased by 22.1%, 22.9%, 9.4% and 19.6% for CPM-P, CPM-N, SVR and LASSO separately. Similar to bootstrapping without replacement, bootstrapping with replacement displayed best accuracies in SVR (average r=0.26), suboptimal result in LASSO (average r=0.22) and similar results in CPM-P (average r=0.18) and CPM-N (average r=0.19). The best prediction accuracies of 13 cognitive measures were illustrated in Figure 3 for bootstrapping without and without replacement.

**Figure 2.**
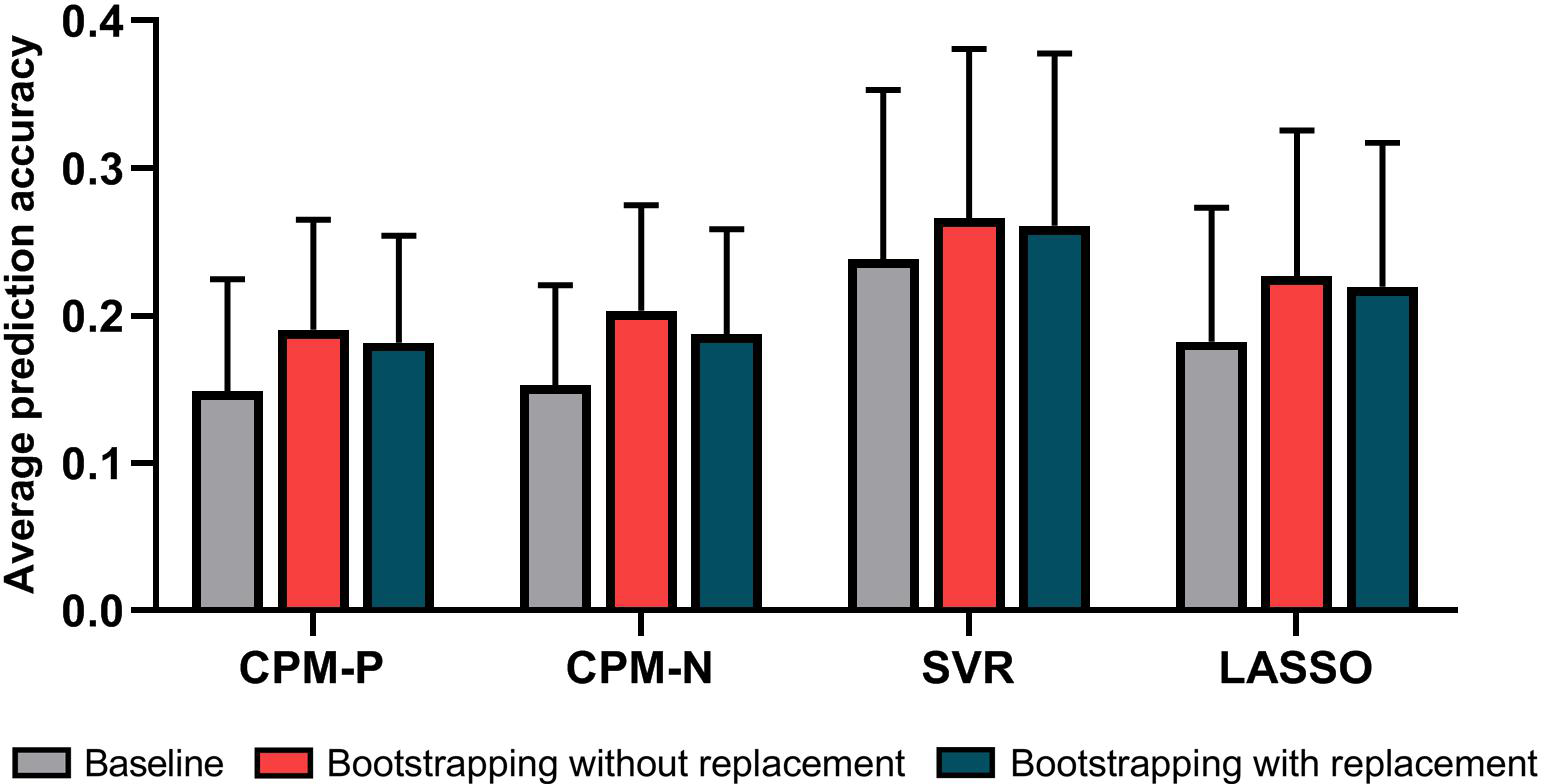
The average best prediction accuracies of bootstrapping methods and baseline method with CPM-P, CPM-N, SVR and LASSO model.

**Figure 3.**
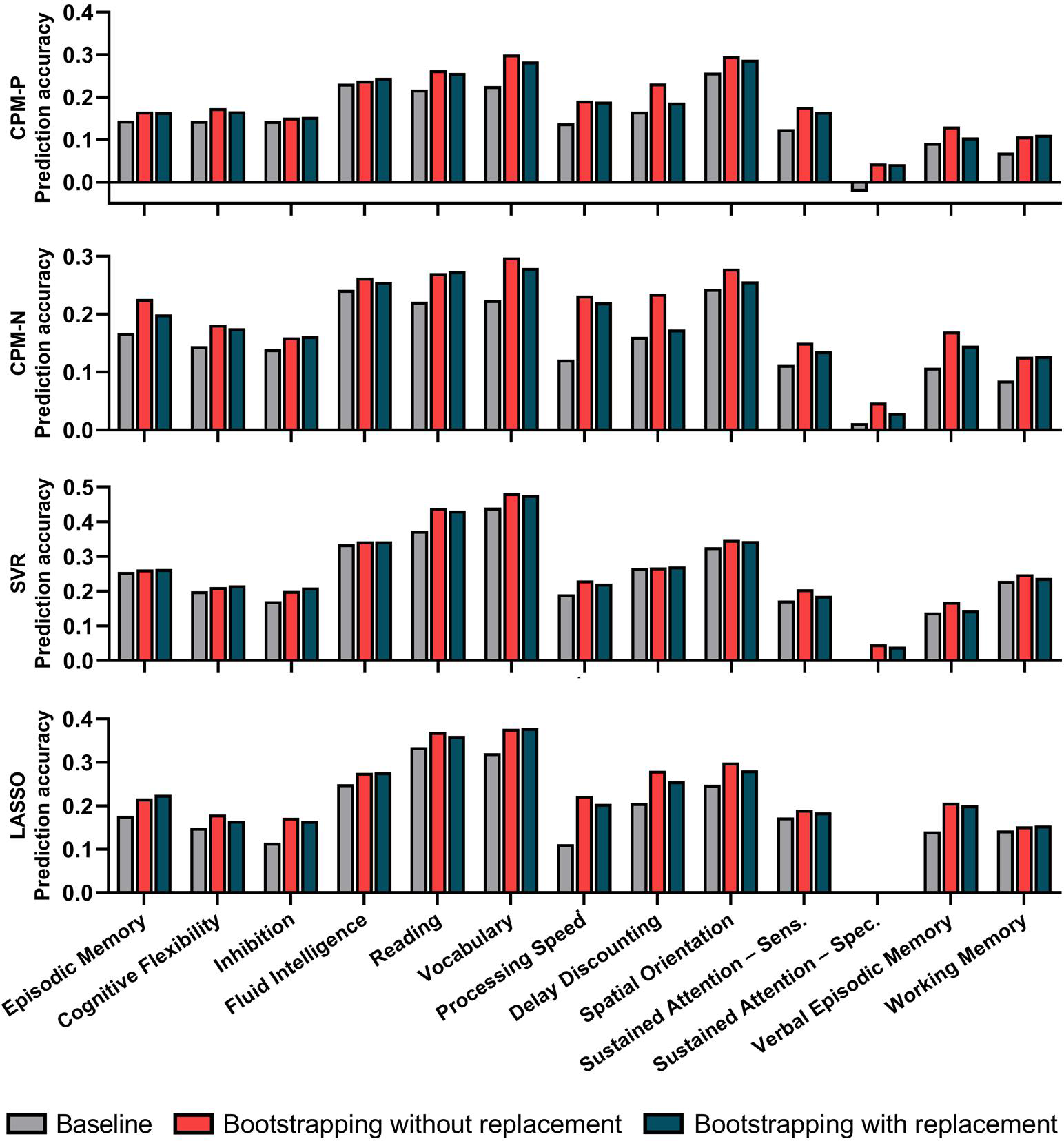
The best prediction performances for 13 cognitive measures with CPM-P, CPM-N, SVR and LASSO.

### 3.2 Bootstrapping methods largely reduced feature dimension

The feature dimension refined by bootstrapping methods decreased enormously compared with the baseline method (Figure 4). Using bootstrapping without replacement, the average feature dimension decreased from 2574 to 209 in CPM-P, from 2648 to 136 in CPM-N, from 5222 to 1359 in SVR and from 5221 to 351 in LASSO. Using bootstrapping with replacement, the average feature dimension decreased to 521 in CPM-P, 188 in CPM-N, 2892 in SVR and 1771 in LASSO. Generally, bootstrapping without replacement reduced more features than bootstrapping with replacement.

**Figure 4.**
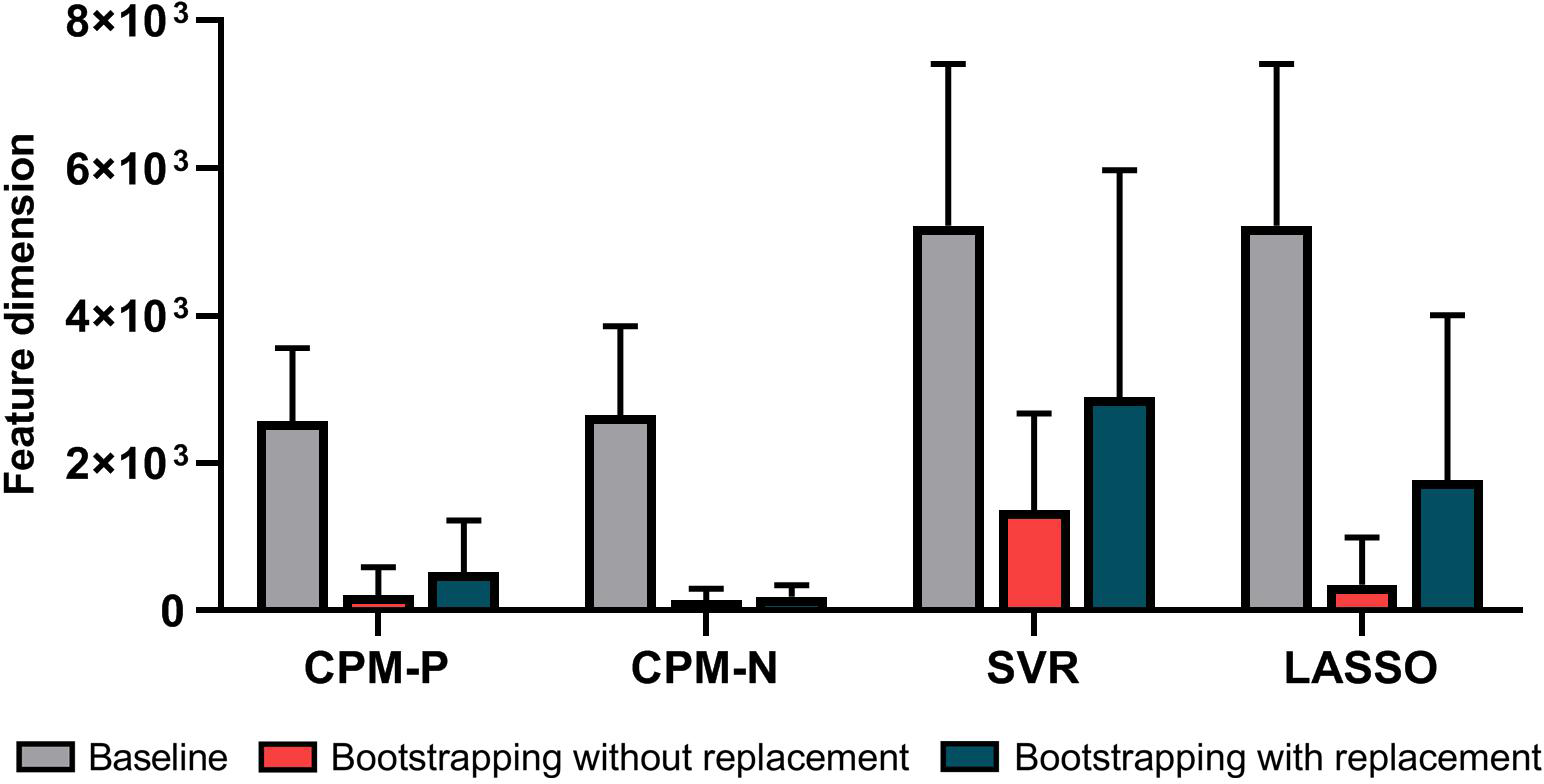
The comparison of feature dimensions between baseline and bootstrapping methods.

### 3.3 Time consumption was dominated by number of bootstrapping

The time consumption of the bootstrapping methods gradually increased with *NB* (figure 5). Under the same *NB*, bootstrapping with replacement consumed more time than bootstrapping without replacement, which demonstrated the efficiency of bootstrapping without replacement. When *NB* exceeded 500, the average consumption time became significantly larger, implying that *NB* was preferably set to be less than 500.

**Figure 5.**
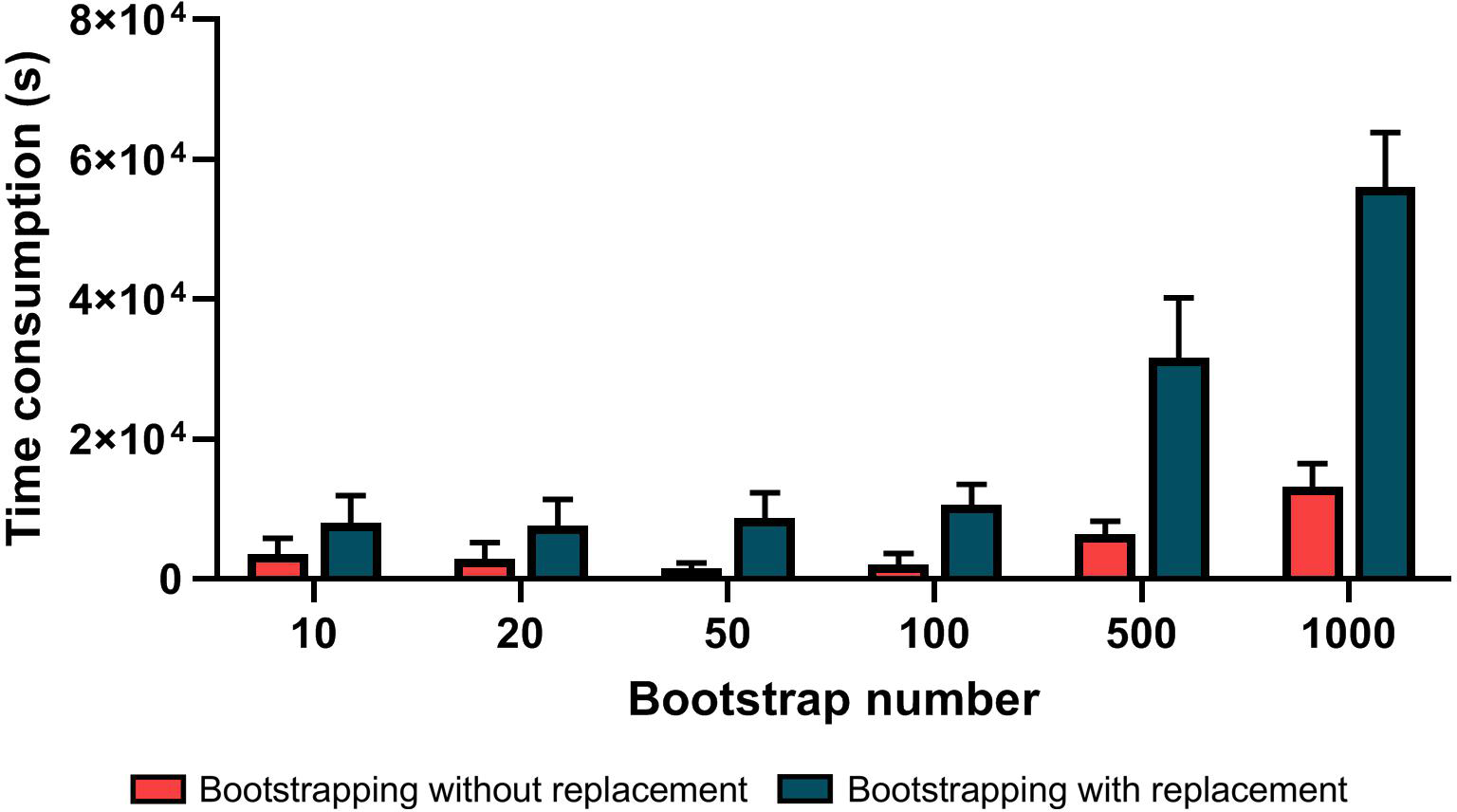
The average time consumption in bootstrapping methods for 13 cognitive traits.

### 3.4 Optimal parameter settings were not constant for different cognitive traits

The parameter sets (i.e., *NB*, *FP* and *BP*) which achieved best prediction performance for each cognitive measure were depicted in Figure 6. Here, a dot in the parameter space referred to the best parameter setting for one cognitive measure. The results showed most of the cognitive scores could be best estimated by small values of *NB*, and 83% and 88% of best performances were respectively achieved with *NB* less than 100 for bootstrapping without and with replacement. As for *FP* and *BP*, they varied in different models for each cognitive trait. For CPM, large *BP* (around 0.8) and moderate *FP* (0.6-0.8) achieved good performances in bootstrapping without replacement, while high *FP* (0.8-1.0) was optimal for bootstrapping with replacement. For SVR, it appeared that there were no obvious rules in *BP* and *FP*. For LASSO, a large *BP* (0.7-0.8) seemingly obtained good performances in bootstrapping without replacement.

**Figure 6.**
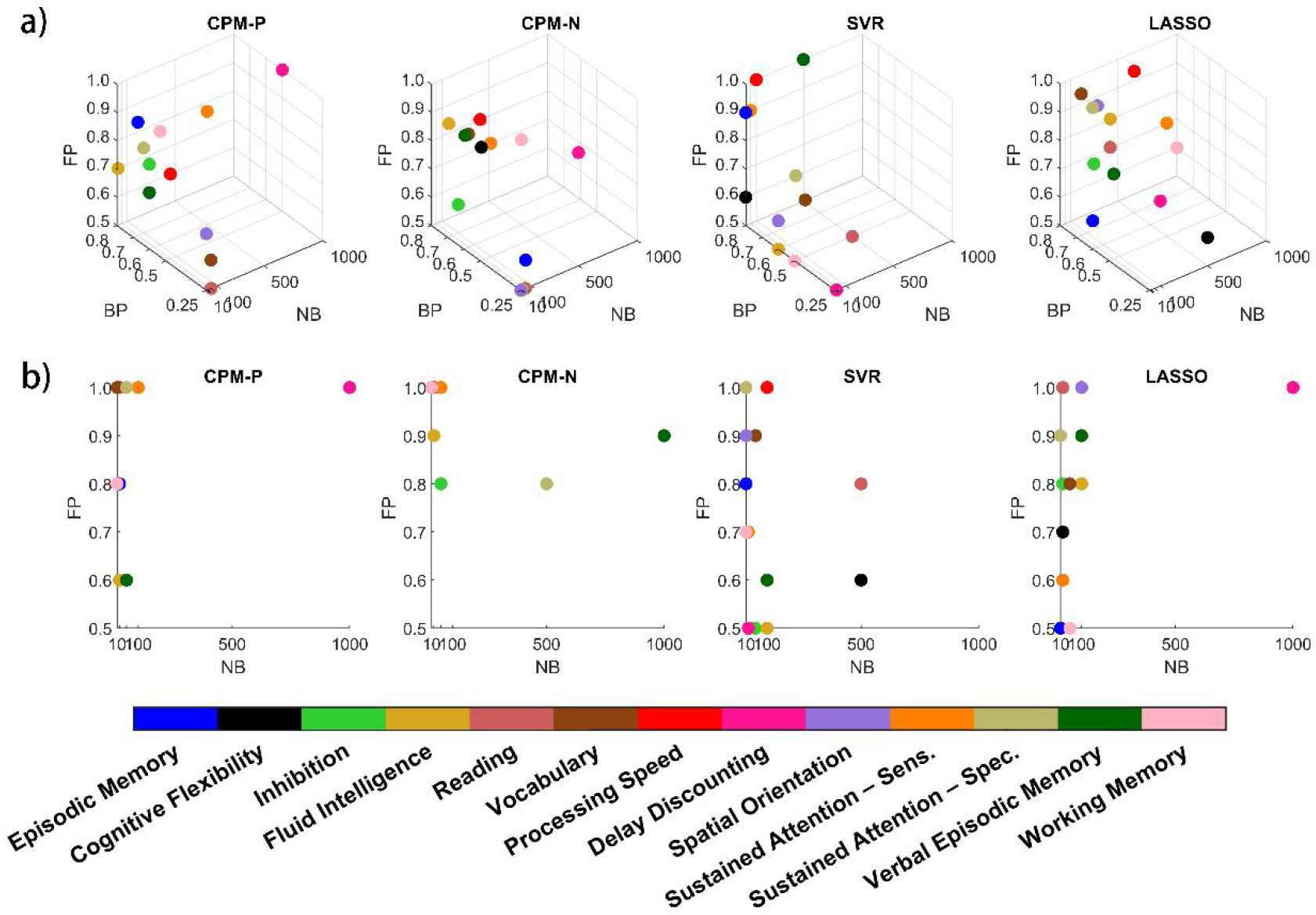
The optimal parameter setting for each cognitive measure in CPM-P, CPM-N, SVR and LASSO when using **a)** bootstrapping without replacement and **b)** bootstrapping with replacement. The colored dots denoted different types of cognitive scores.

## 4 DISCUSSION

In order to improve the performance of RSFC prediction, this study developed bootstrapping based feature selection framework for RSFC prediction of cognitive traits. Using a large sample from HCP dataset, the prediction of cognitive traits with bootstrapping methods significantly outperformed the baseline CPM, SVR and LASSO models, and the dimension of selected features decreased dramatically using the bootstrapping methods. In addition, bootstrapping without replacement displayed superior improvements and higher running efficiency in contrast to bootstrapping with replacement.

### 4.1 Bootstrapping enhanced the RSFC-phenotype associations

Modeling cognitive traits with RSFC is a persistent pursuit in modern cognitive neuroscience studies (Petersen et al., 2015; Poldrack et al., 2015). A previous study shows that the diversity of individual cognition can be represented by the inter-subject variations in the FC patterns (Mueller et al., 2013). However, the FC patterns may be noisy and unreliable (Braga et al., 2017; Gordon et al., 2017; Gratton et al., 2018; Mueller et al., 2013), thus confound the prediction precision. In recent years, improving RSFC-phenotype association is a major concern in the domain, and several methodologies have been proposed from different perspectives. The first way is to use individual-specific RSFC patterns to predict the behavioral traits. In Qin et al. (2019) and Kashyap et al. (2019), they decompose RSFC (or timecourse) to extract individual-specific RSFC (or timecourse) with different methods, which obtained obvious improvements in prediction. A second way is to combine various kinds of information with RSFC to enhance the prediction, such as task-fMRI based FC (Elliott et al., 2019; Gao et al., 2019; Xiao et al., 2019) and dynamic FC (Liegeois et al., 2019; Lim et al., 2018; Park et al., 2018), which can provide complementary information to the conventional FC. A third way is to decrease the influence of the possible noise in rs-fMRI signal, for instance, global signal regression (Li et al., 2019) and motion artifact correction (Nielsen et al., 2019) have been reported to advance the RSFC-behavior prediction. The last way is to use the bagging strategy (Breiman, 1996) to improve the prediction with RSFC (Jollans et al., 2019).

Our proposed method provides another perspective to promote the RSFC-phonotype prediction, and displays several advantages. First, the method does not need any collection of additional data (e.g. task fMRI) or any other processing (e.g. computation of dynamic FC and FC decomposition), so it is an easy-to-use and effective method. Second, the current method can purify the selected feature sets, making the prediction model more succinct and interpretable. It is known that reducing feature dimension is always beneficial to attenuate overfitting in neuroimaging data (Jollans et al., 2019). At last, previous studies (Alonso-Atienza et al., 2012; Ditzler et al., 2015) used bootstrapping or bagging in a computation expensive manner, which requires the model construction and optimization in each bootstrapping resample dataset. This may lead to an unacceptable running time for the heuristic searching of optimal parameter setting in large neuroimaging dataset, and only have to use constant parameter setting to conduct such methods.

As we know, good stability of biomarkers is thought as important as high classification performances (He et al., 2010). Currently, the importance of the stability of the rs-fMRI signal (e.g. test-retest reliability) and the stability of selected biomarkers across sites (e.g. multi-center dataset) have been widely accepted (Marchitelli et al., 2016; Mueller et al., 2015; Noble, Scheinost, et al., 2017; Noble, Spann, et al., 2017), however, there is seldom study special focusing on the stability of selected biomarkers across subsamples in one dataset. Indeed, heterogeneity exists in nearly all human neuroimaging datasets regardless of whether the samples are healthy or not, which may therefore result in instability in feature sets derived from subsamples and further weaken the prediction performance. It is necessary to eliminate the heterogeneity in the dataset and to obtain stable representations standing for neural activities. Figure 4 clearly shows that the amounts of false correlated feature sets are large, implying that it is very beneficial to take into account the stability of feature selection for machine learning studies with neuroimaging dataset. Our study demonstrated that the proposed bootstrapping-based framework is an effective way to eliminate the heterogeneity in the dataset and to obtain feature sets with good stability.

### 4.2 Bootstrapping with replacement vs. bootstrapping without replacement

Bootstrapping with and without replacement were compared in the study. Though bootstrapping with replacement was referred more frequently than bootstrapping without replacement, it was reported not to guarantee reliable results sometimes. In Strobl et al. (2007), feature selection in random forest was biased by bootstrapping with replacement, while importance measures of predictors could be accessed more reliably by bootstrapping without replacement. This phenomenon was not hard to explain, bootstrapping with replacement ensured the resample size as same as the original dataset, so the replication samples might introduce possible bias into the final feature selection, which was especially obvious in a finite dataset (e.g. neuroimaging data). For bootstrapping without replacement, the resamples was a subset of original dataset, so they could be regarded as sampling from the same population, and properties of statistical test in resamples were constant with the original data (Rospleszcz et al., 2016). Moreover, bootstrapping without replacement obtained greater prediction improvements, reduced more features that were redundant and ran more efficiently than bootstrapping with replacement. Hence, we recommended bootstrapping without replacement as a feature selection technique to draw reliable predictors.

### 4.3 Comparison of CPM, LASSO and SVR

For the prediction performance, CPM achieved the worst performance compared with SVR and LASSO. There were three possible aspects accounted for it: 1. CPM is a simple linear regression model, while SVR and LASSO are models with L1 and L2-norm regularization, which may therefore obtain better prediction performances and also increase the model generalization. 2. The correlated feature sets were respectively divided into positive and negative feature sets in CPM, while SVR and LASSO used their combination, so such division may weaken the CPM performance significantly. 3. CPM assigned same weight to all features, which might underestimate key predictors and overestimate weak predictors. Notably, CPM achieved the largest improvement in prediction and the most obvious reduction of features using bootstrapping methods, suggesting future studies with CPM should consider the bootstrapping methods.

LASSO achieved suboptimal prediction performance in three models and it ensured sparseness by randomly selecting one feature from correlated features. LASSO was shown to perform well in challenging situation like potential features size exceeded the sample size (Abraham et al., 2017; Bunea et al., 2007a; Bunea et al., 2007b; Zhang et al., 2008). However, when confronting the condition that the feature number was remarkably larger than the sample size, it might meet such a problem: LASSO could only retain no more than N−1 features (N is the sample size), thus some informative features might be discarded in the model (Efron et al., 2004; Ryali et al., 2012).

SVR displayed the best performance among three models. Compared with LASSO, SVR performed slightly better and the similar trends were also revealed by Cui et al. (2018) and Dadi et al. (2019) on RSFC predictions. However, efficiency was a crucial problem for high-dimensional data while SVR was tremendously expensive in computation for model fitting (Cui et al., 2018; Shen et al., 2017).

### 4.4 Optimal selection of parameter setting

In bootstrapping with and without replacement, most best performances were achieved with *NB* less than 100, implying that bootstrapping did not require a mass of resampling times to acquire stable features, so we recommended small *NB* (<100) for bootstrapping methods used in CPM, SVR and LASSO models. *PF* indicated the stability of the feature in the resampling subset, thus specifying an appropriate threshold for *FP* was a key setting. In Bunea et al. (2011), the *FP* threshold was set to a single value (50%) in LASSO in order to not miss any possibly relevant features, but our study found the optimal *FP* threshold varied for different cognitive traits, indicating that constant threshold setting could not guarantee the best performance in LASSO. For CPM, the optimal *PF* threshold was moderate values (0.6-0.8) but not the very high values (0.8-1) in bootstrapping without replacement, which reflected that outliers and clustered points might also exist in the bootstrapping resamples, therefore, *FP* threshold should not be set with very high values in CPM. *BP* stood for the resample percentage from the original dataset, and our results discovered that moderate values (0.7-0.8) were suitable for CPM and LASSO in bootstrapping without replacement. For SVR, there were no significant rules for all parameter settings, so the best parameter settings should be searched for each prediction target.

### 4.5 Other methodological considerations

Group ICA has been found to be a better way to define the functional brain nodes than the atlas-based method for its better prediction accuracy (Dadi et al., 2019), hence, group ICA parcellation was adopted to calculate the corresponding FC in the study. There were various brain parcellations derived from group ICA provided in HCP dataset, i.e., [15, 25, 50, 100, 200, 300]. To have direct comparisons among different parcellations, the baseline CPM, SVR and LASSO methods were used to predict the cognitive functions with FC matrix under different parcellations. The results were illustrated in Figure 7, and the main tendency was that the prediction accuracy increased with dimension except LASSO, and grown slowly from 100 to 300 independent components, and similar trend was also revealed in other works (Abraham et al., 2017; Dadi et al., 2019). To investigate the bootstrapping methods on high-dimensional data, the analysis was based on 300 independent components. Besides, significant threshold p-value was an important parameter for correlation analysis. In order to conserve as much as possible features, the p-value was set at 0.05 rather than other more stringent value in this study. In addition, concerning different head motion correction strategies and global signal regression were still controversial in functional connectivity studies, and we just focused on the bootstrapping related improvement, so we did not compare any other preprocessing steps on the preprocessed HCP data.

**Figure 7.**
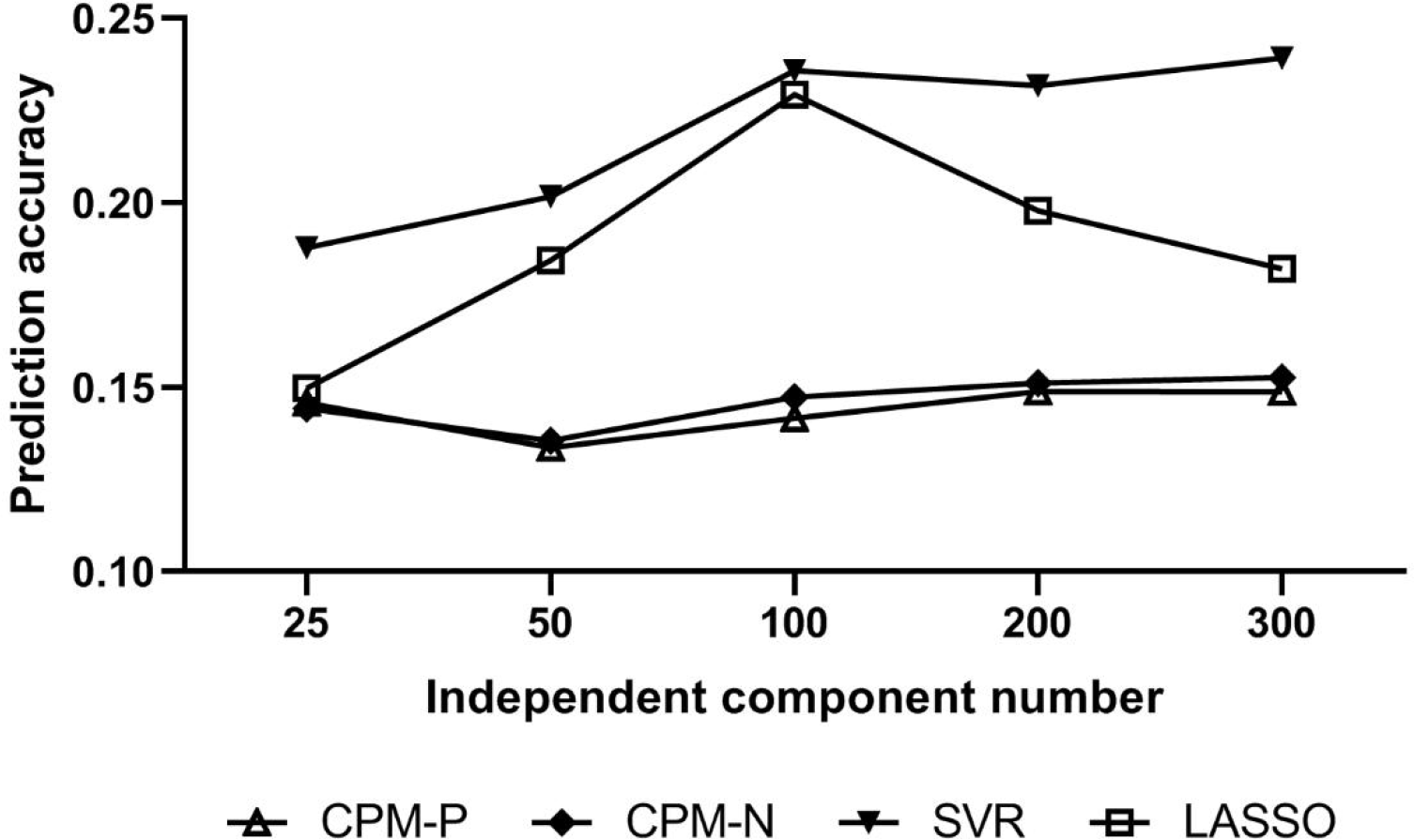
The prediction accuracy of different number of independent components. The result of 15 parcels was not shown since no significant features were selected for some cognitive scores.

### 4.6 Limitation and future work

The bootstrapping methods were more expensive in computation compared with the baseline method, which could be speeded up via parallel computing with multiple CPUs, making the bootstrapping methods feasible in practical use. An additional disadvantage was that the optimal parameter settings varied in different cognitive phenotype predictions, so a heuristic search was needed for new prediction target. Although the bootstrapping method could improve the RSFC-phenotype association, the prediction accuracy was still not high enough, and future work should combine other improvement ways together to make the RSFC-based prediction applicable for clinical requirement.

## 5 CONCLUSION

This study presented a bootstrapping-based feature selection framework and was applied to CPM, SVR and LASSO models, the results demonstrated the bootstrapping method could not only improve the RSFC-phenotype association but also purify the selected features into a lower dimension. Future RSFC prediction works were highly recommended to use the bootstrapping method.

## ACKNOWLEDGMENTS

We are grateful to Xiangyu Ma from School of Biomedical Engineering, Capital Medical University, Beijing for valuable comments for the manuscript. This work was supported by the Beijing Municipal National Science Foundation (4122018).

## DATA AVAILABILITY STATEMENT

Data were provided by the Human Connectome Project and WU-Minn Consortium (Principal Investigators: David Van Essen and Kamil Ugurbil; 1U54MH091657) funded by 16 NIH Institutes and Centers that support the NIH Blueprint for Neuroscience Research; and by the McDonnell Center for Systems Neuroscience at Washington University. Code for this work is freely available at the GitHub repository (https://github.com/Lijiang-Wei/bootstrapping.git).

